# Functional pathways for metabolic network-based data analysis: the MetPath algorithm

**DOI:** 10.1101/202226

**Authors:** Gianluca Mattei, Daniel C. Zielinski, Zhuohui Gan, Matteo Ramazzotti, Bernhard O. Palsson

**Affiliations:** Department of Bioengineering, University of California, San Diego, La Jolla CA 92093-0412; Department of Experimental and Clinical Biomedical Sciences, University of Florence, Florence, Italy.

## Abstract

Analyzing biological data using pathways helps identify trends in data tied to the function of a network. A large number of pathway-based analysis tools have been developed toward this goal. These pathways are often manually curated and thus associations are subject to the biases of the curator. A potentially attractive alternative is to define pathways based on the inherent functionality and connectivity of the network itself. Within metabolism, functionality is defined by the production and consumption of metabolites, and connectivity by metabolites participating in reactions through common enzymes. In this work, we present an algorithm, termed MetPath, that calculates pathways for production and consumption of metabolites. We show how these pathways have attractive properties, such as the ability to integrate multiple data types and weight contribution of genes within the pathway by their functional contribution to metabolite production/consumption. Pathways calculated in this manner are condition-specific and thus are custom tailored to the system of interest, in contrast to curated pathways. We find that these pathways predict gene expression correlation better compared to manually-curated pathways. Additionally, we find that these pathways can be used to understand gene expression changes between growth conditions and between cell types. This work serves to better understand the functional pathway structure underlying cell metabolism and helps to enable systems analyses of high-throughput data.

## Introduction

In the modern era of biology, high-throughput molecular data such as genome-scale gene expression measurements are ubiquitous. However, this abundance of data comes with the challenge of extracting knowledge from the data, which is largely responsible for the rise of systems biology. As cells are fundamentally composed of networks of interacting molecules, the data analysis challenge comes down to obtaining an understanding of how each of the data points interacts in the context of the underlying biochemical network. Systems biology methods of data analysis can be roughly grouped into network topology, ontology, and pathway-based methods. Within metabolism specifically, pathway analysis has played a particularly important role(Khatri, Sirota, and Butte 2012), as connections between enzymes have clear functional objectives in conversion of molecules to energy and biomass. However, identifying pathway structures that are best used to interpret high-throughput data remains an open challenge.

Metabolic pathways have historically been defined manually based on an intuitive understanding of the relationship and function of particular sets of enzymes. Classical examples such as glycolysis and the TCA cycle appear in textbooks and databases with well-established structure and content(Kanehisa et al. 2017; Subramanian et al. 2005). However, canonically-defined pathways are not necessarily the most optimal pathways for the purpose of understanding organism function, and they do not generally take into account the myriad variations of pathways that exist across the phylogenetic tree nor the condition-specific use of pathways. As an alternative to manually-defined pathways, a number of methods have been developed to algorithmically calculated pathways from the metabolic network structure directly(Bordbar et al. 2014). These algorithmic methods often have a strict numerical objective, such as the extreme pathways that are a non-negative basis for the nullspace of the stoichiometric matrix of the metabolic network(Wiback, Mahadevan, and Palsson 2003). Still, these numerical objectives may not be the most practical nor the most effective pathways to interpret high-throughput data.

The function of a metabolic network can reasonably be defined as the production and degradation of metabolites. To more rigorously define the machinery that accomplishes these functions in a systems context, we can define pathways, or sequential sets of enzymes, that are involved in the production and/or degradation of each metabolite under a defined metabolic flux state. An intuitive definition of a metabolic pathway thus is a series of consecutive reactions that result in the production of a metabolite or consumption of a metabolite. This definition inherently assumes that the flux directions through the network are defined, but in fact these are not fixed across conditions. A set of reactions may be involved in consumption of a metabolite under one condition, and production of the metabolite under another condition, for example the relationship between reactions in glycolysis and glucose under glycolytic compared to gluconeogenic conditions. Thus, the functional interpretation of changes in enzyme levels depends on the flux directions in the network. Furthermore, the relative importance of enzyme changes depends on the contribution of a pathway to a metabolic function. As an example, the pathways of glycolysis and glycogen synthesis both consume glucose. However, glycolysis operates at a rate approximately 100x that of glycogen synthesis. Thus, a 10% increase in expression of glycolytic enzymes would greatly outweigh a 10% decrease in glycogen synthesis, resulting in an estimated net decrease in the metabolic capability to consume glucose. We can use this type of analysis to examine how gene expression has changed in these pathways in order to to identify coordinated expression shifts that serve specific metabolic functions in terms of increased or decreased capacity for production or degradation of specific metabolites.

In this work, we develop a constraint-based modeling method, termed MetPath, to calculate condition-specific production and consumption pathways for the purpose of differential analysis of gene expression data. Using metabolic modeling, we can calculate metabolic pathways for specific flux conditions, defining weighted, context-specific pathways defined in terms of metabolic functions, i.e. production/consumption of specific metabolites. We compare this method qualitatively and quantitatively to existing pathway databases, most notably KEGG, and find that MetPath pathways show higher intergene correlation within pathways across conditions in *E. coli* K12 MG1655. We examine the performance of MetPath on two case studies in *E. coli*, looking at the tryptophan pathway during aerobic growth on glucose with and without tryptophan supplementation as well as glucose growth in the aerobic-anaerobic shift. Finally, we look at the ability of MetPath pathways to interpret cell-specific differences in gene expression, examining specifically neurotransmitter pathway expression in human neural cell subtypes based on single cell transcriptomics data.

## Results

### Calculation of condition-specific pathways for production and consumption of metabolites

The MetPath computational workflow to analyze the change in metabolic production and degradation pathways in the network is as follows (Figure 1). First, we define an estimated metabolic state based on established metabolite uptakes and energy production estimates. We then define production and degradation pathways for each metabolite in the network using the network structure and constraint-based pathway definition algorithms(Orth, Thiele, and Palsson 2010). Finally, we create an aggregate perturbation score for each production and degradation pathway based on the fold change of significantly changed metabolic genes within each pathway. Perturbed pathways thus represent network-integrated gene expression changes in production or degradation potential of specific metabolites.

**Figure 1:**
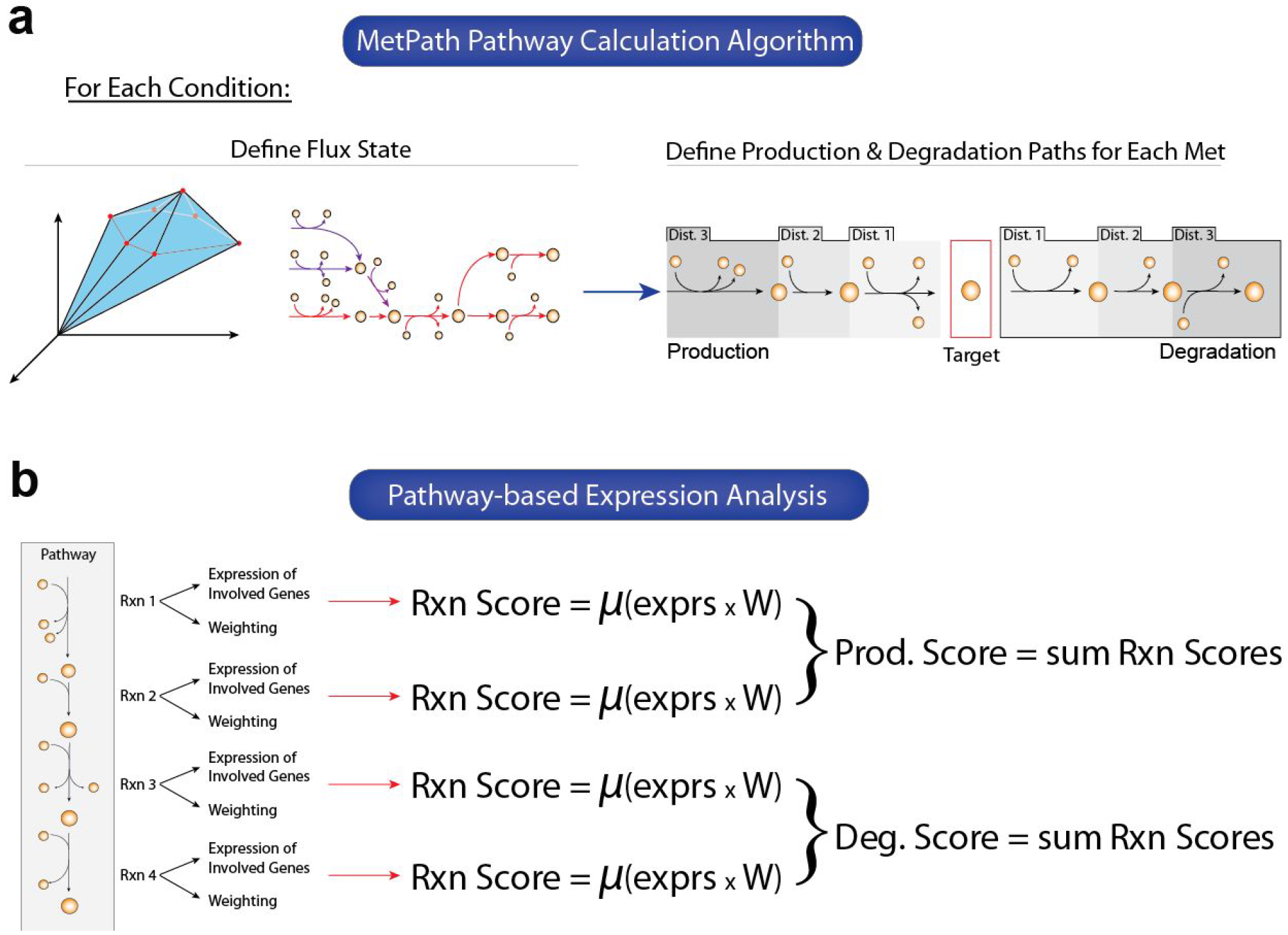
Overview of the MetPath algorithm. **a)** MetPath is a constraint-based modeling method to calculate pathways for the production and consumption of each metabolite in the network. The algorithm begins by calculating a flux state for a condition of interest. Then, for each metabolite, a subnetwork around the metabolite that is active based on the flux state for the condition is extracted. This subnetwork is then broken down into production pathways and consumption pathways, weighted by their flux contribution, using elementary modes. **b)** To interpret differential gene expression data using MetPath pathways, a reaction score is calculated for each reaction in the pathway as the multiplication of the differential gene expression for genes catalyzing the reaction with the weighting on that reaction within the pathway. These reaction scores are summed and divided by the number of reactions to obtain a pathway score. A value of 1 indicates unchanged expression for the pathway, greater than one indicates an up-regulation of genes in the pathway, and less than one indicates a down-regulation of genes in the pathway.

As the definition of a metabolic state depends on the condition of interest, we defined a set of standard conditions that may be of interest to users and calculated MetPath pathways for each of these. We utilized the iJO1366 *E. coli* genome-scale metabolic network(Orth et al. 2011) and calculate flux states for 40 conditions, altering carbon and nitrogen source as well as terminal electron acceptor. We combined the MetPath pathways for each metabolite in the network for these conditions into a single database, combining pathways that were shared between conditions (MCC between the pathways greater than a cutoff) and leaving dissimilar pathways separate. The resulting pathway database consists of a set of condition-specific production and consumption pathways for each metabolite in the network. This database serves as a basis for pathway-based analysis of gene expression data across diverse conditions.

### Comparison of pathway behavior to other pathway databases

To assess the ability of the MetPath *E. coli* pathway database to interpret differential gene expression, we gathered 213 gene expression samples from *E. coli* K12 MG1655 grown under various conditions and genetic perturbations. We mapped this gene expression data onto the pathway database and examined co-expression of genes within pathways. We calculated the correlation of genes within pathways compared to genes that do not share pathways (Figure 2A) and compared to pathways extracted from the KEGG database. We found that genes within MetPath pathways are substantially more correlated. This correlation is dependent upon the pre-defined length of MetPath pathway, with shorter distances associated with higher correlations. This indicates that co-expression of genes along metabolic pathways tends to be highly colocalized. As KEGG pathways tend to be significantly longer than MetPath pathways, this colocalization bias leads to a lower total co-expression of genes in KEGG pathways. Additionally, we found that higher expressed genes tended to be more co-expressed within pathways (Figure 2B). This could indicate either tighter gene expression regulation of highly expressed genes, or clearer correlation signal in highly expressed genes compared with low expressed genes due to noise associated with measuring the latter.

**Figure 2:**
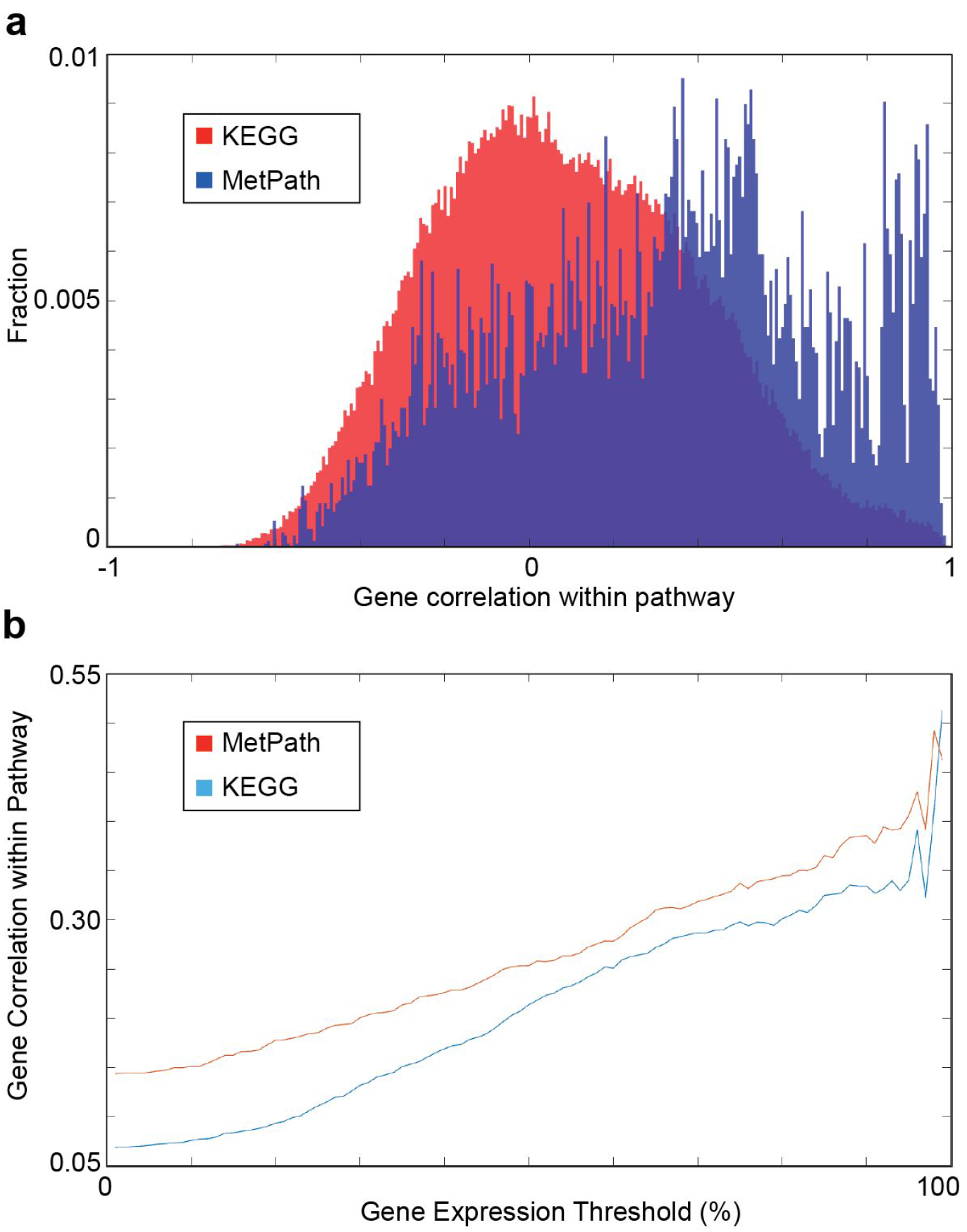
Comparison of MetPath and KEGG pathways in predicting correlated gene expression correlation. **a)** Histograms of the correlation of gene expression within pathways for KEGG (red) and MetPath (blue) pathways. Expression data was obtained from 213 samples under various conditions in E. coli K12 MG1655. **b)** Gene correlation within pathways as a function of expression level of the genes. Highly expressed genes show greater correlation with each other within pathways that low expressed genes.

### MetPath pathways reveal coordinated expression changes with shifts in environment

We then were interested in determining whether MetPath pathways can identify functional differences between metabolic states based on gene expression data. Again utilizing available gene expression data for *E. coli*, we examined two comparisons. First, we looked at the MetPath production and consumption pathways for tryptophan for *E. coli* grown aerobic on glucose with and without tryptophan supplementation (Figure 3A). We found that due to the change in underlying metabolic flux state calculated with flux balance analysis for each condition, the MetPath pathways differ substantially between the glucose only case, where tryptophan must be synthesized *de novo*, and the tryptophan supplemented case. This condition change is associated with both a clear expression change that is observed through the MetPath pathway scores as well as a known shift in activity of the transcription factor regulating this pathway. Second, we examined central energy and oxidative metabolism during the aerobic-anaerobic shift in *E. coli.* Looking specifically at the pyruvate pathway, a key branch point in the oxidative/glycolytic shift (Figure 3B), we once again see a coordinated multi-gene response along MetPath pathways. This indicates that a clear expression signature exists that can be mapped onto the metabolic network through MetPath pathways to obtain an integrated signature with potentially greater statistical power than single gene-based expression analyses.

**Figure 3:**
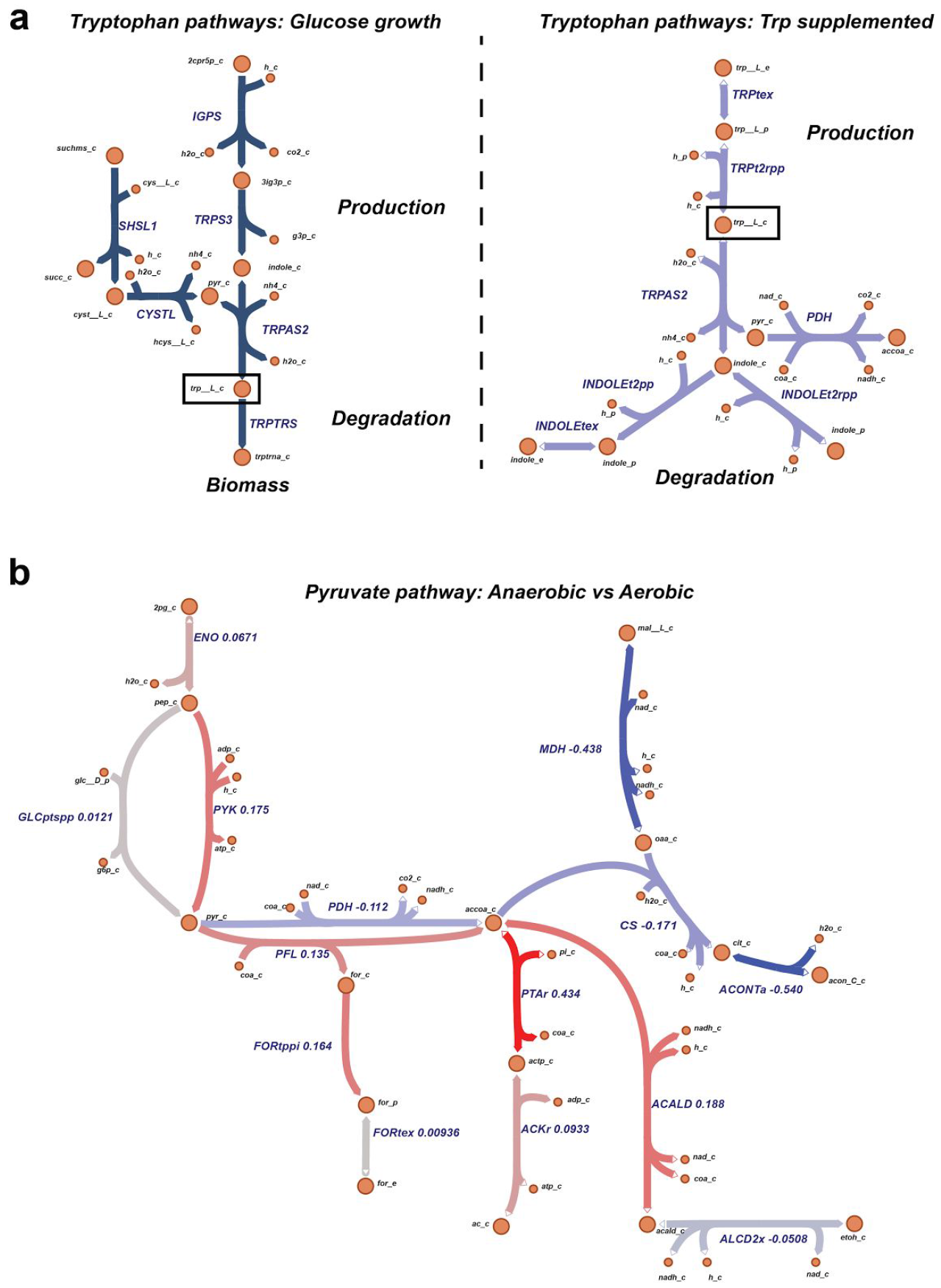
MetPath pathways highlight metabolic shifts due to growth condition. **a)** MetPath pathways under two conditions, glucose growth and tryptophan supplementation, reveal differential pathway definition of tryptophan production and consumption in a condition-specific manner. **b)** Differential expression between aerobic growth and anaerobic growth of *E. coli* revealed by MetPath scores for pyruvate.

### MetPath pathways classify neural cells by neurotransmitter pathway expression

Finally, we wanted to determine whether MetPath pathways could identify functional metabolic differences across entirely different cell types using their gene expression alone. To this end, we utilized single cell gene expression data from a set of 33 cell subtypes identified from human brain samples (Lake et al. 2016). Given that the samples originated from the human brain, we were interested specifically in whether MetPath could identify differential use of metabolic pathways associated with neurotransmitters in these cell subtypes. We collected pathways for a representative set of neurotransmitters and mapped expression data for the neural cell subtypes onto these pathways in the global human metabolic network reconstruction Recon 1(Duarte et al. 2007) (Figure 4A). Encouragingly, we observed clear differentiation of cell subtypes based on expression of neurotransmitter pathways. These neurotransmitters matched canonical use within particular neural cell subtypes, such as the association of GABA with inhibitory neurons and glutamate with excitatory neurons. Additionally, we identified unusual neurotransmitter use among particular subtypes of neurons, such as an up-regulation of the NO synthesis pathway among particular subtypes of inhibitory neurons. Again with the goal of comparing integrated pathway analysis with single gene analysis, we extracted the glutamate pathway as a case study (Figure 4B). We compared expression of the glutamate pathway within excitatory neurons where this pathway is canonically activate and endothelial cells where its role is unclear. As with *E. coli*, we observed that there is a coordinated multi-gene signature of up-regulation of glutamate production in the excitatory neuron. This signature includes an up-regulation of glutamate production and secretion genes and down-regulation of the primary glutamate degradation enzyme Glutamate Dehydrogenase was observed, consistent with the use of glutamate as an excitatory neurotransmitter. Thus, it appears that the integrated analysis using MetPath pathways reveals additional coordination that would be more difficult to see based on single gene analysis.

**Figure 4:**
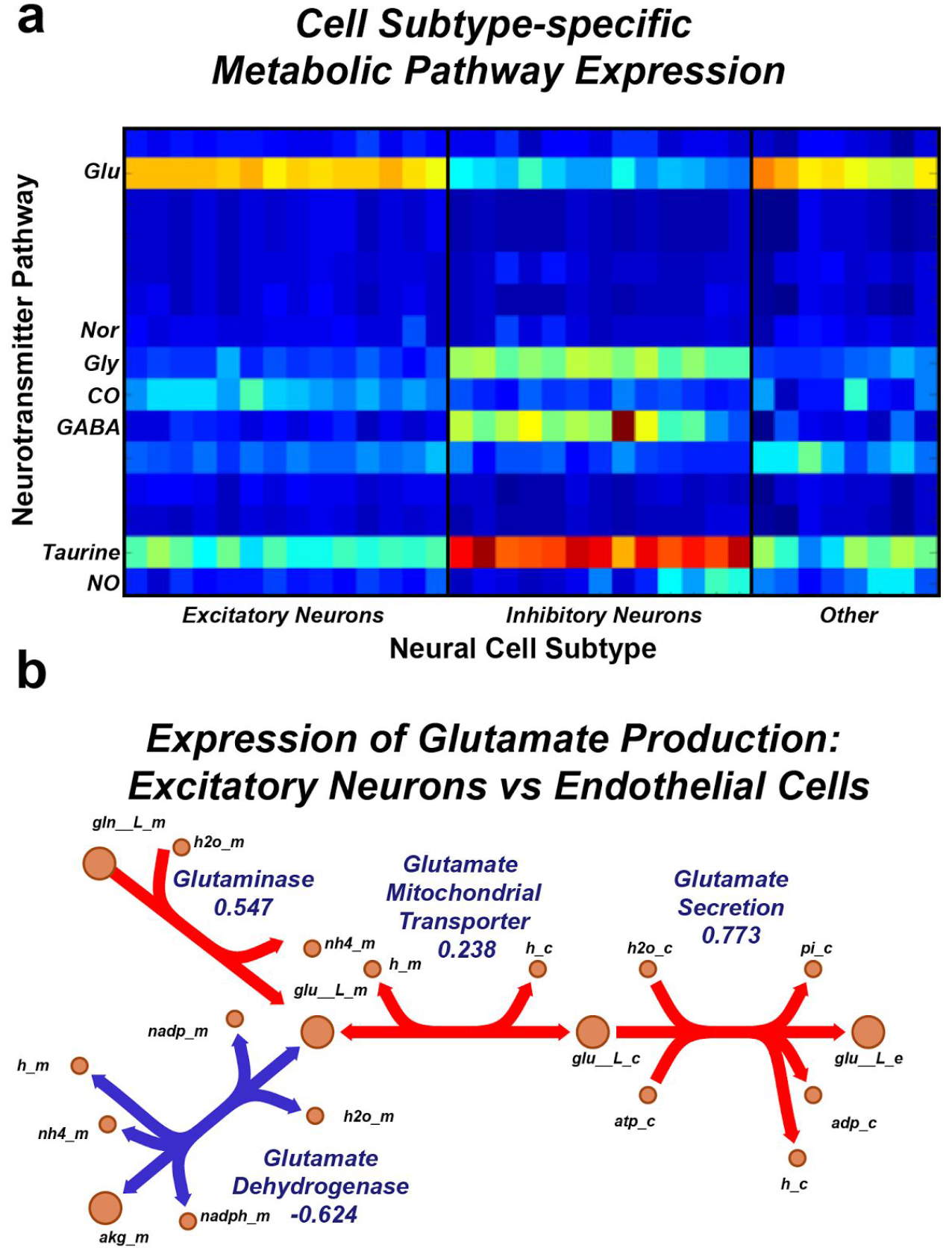
MetPath reveals cell-specific expression of neurotransmitter use in single cell neural gene expression data. **a)** Single cell gene expression data mapped to metabolic pathways for neurotransmitter production for a representative set of neurotransmitters. Subtypes of neural cells revealed differential expression of neurotransmitters consistent with canonical neurotransmitter use. **b)** MetPath scores for glutamate production in excitatory neural cells compared with endothelial cells. A coordinated up-regulation of glutamate production and secretion genes and down-regulation of the primary glutamate degradation enzyme Glutamate Dehydrogenase was observed, consistent with the use of glutamate as an excitatory neurotransmitter.

## Discussion

In this work, we developed a constraint-based modeling method, termed MetPath, to calculate production and consumption pathways for the purpose of differential analysis of gene expression data. We compared this method qualitatively and quantitatively to existing pathway databases, most notably KEGG, and found that MetPath pathways showed higher intergene correlation within pathways across conditions in *E. coli* K12 MG1655. We examined the performance of MetPath on two case studies in *E. coli*, looking at the tryptophan pathway during aerobic growth on glucose with and without tryptophan supplementation as well as glucose growth in the aerobic-anaerobic shift. Finally, we looked at the ability of MetPath pathways to interpret cell-specific differences in gene expression, examining specifically neurotransmitter pathway expression in human neural cell subtypes based on single cell transcriptomics data.

The basis for using a metabolic pathway to understand the purpose of differential gene expression is rooted in the assumption that the expression difference is associated with a change in flux through the metabolic pathway. The link between gene expression and metabolic flux, which is the variable of greatest interest, is known to be indirect at best (Chubukov et al. 2013). mRNA and enzyme levels are known to have only modest correlation(Fendt et al. 2010), and the flux catalyzed per unit of enzyme depends also on metabolite levels, which can change between conditions(Bennett et al. 2009). Thus, rather than attempting to estimate differential flux levels directly using gene expression changes, we decided to ask the more addressable question of how gene expression changes have made different ‘functions’ of the metabolic network more or less difficult, on the assumption that mRNA changes and enzyme changes are positively correlated (Greenbaum et al. 2003).

We examined the performance of MetPath pathways related to KEGG pathways in in predicting gene correlation primarily with the goal of displaying the qualitative difference between the behavior of the pathways. This study was not intended to be a rigorous comparison of various pathway databases and pathway algorithms to determine the best performing set of pathways. Others have conducted such analyses(Bordbar et al. 2014), and sufficient increases in available validated data have not been made to warrant revisiting this effort. However, the result that expression among genes was more correlated within MetPath pathways than among KEGG pathways across conditions lends credibility to the hypothesis that the localization of these pathways around production and consumption of individual metabolites, as well as the condition-specific nature of the pathways, may yield some tangible benefits when performing pathway-based data analyses.

To provide real case studies demonstrating the utility of MetPath pathways in understanding the functional significance of gene expression differences, we looked at gene expression from *E. coli* grown under different conditions as well as gene expression from different cell subtypes in the human brain. In each case, we found that highly perturbed pathways were directly tied to the functional difference between conditions or cell type. A similar result may be obtained by looking at expression of individual genes in these cases. However, we observed a coordinated gene expression different among several genes in pathways in each case. Thus, statistical testing around this integrated gene expression change may yield additional power compared to non-pathway-based analyses.

This work may be viewed as one more study in an increasingly long line of pathway definitions for understanding the structure of metabolic networks. The benefits of the MetPath algorithm are that these pathways are 1) automatically calculated, 2) intuitively defined, 3) condition specific, and 4) numerically tractable. We believe that this particular set of traits makes these pathways useful compared to other commonly used pathway sets, and thus this algorithm may be broadly applicable to new studies and organisms.

## Methods

### Overview of MetPath pathway calculation and differential gene expression analysis

We define the functions of a metabolic network as the production and degradation of metabolites. To more rigorously define the machinery that accomplishes these functions in a systems context, we can define pathways, or sequential sets of enzymes, that are involved in the production and/or degradation of each metabolite under a defined metabolic flux state. We can then examine how gene expression has changed in these pathways to identify coordinated expression shifts that serve specific metabolic functions in terms of increased or decreased capacity for production or degradation of specific metabolites.

### Calculation of a condition-specific flux state

To calculate state-specific production and degradation pathways, we first calculate the estimated metabolic state. We solve a flux balance analysis problem on the metabolic model constrained by estimate metabolite uptakes. The flux state is calculated by minimizing the total length of the flux vector subject to the previous constraints, to represent the principle that the cell will try to achieve metabolic function using as little enzyme expenditure as possible to minimize precursor costs. The purpose of the estimated flux state is not to have fully quantitatively accurate flux values for each reaction, but rather to identify likely reaction directions and relative pathway flux values given established literature on aspects such as metabolite synthesis versus *de novo* uptake and relative energy production between glycolysis and beta oxidation. These relative weightings and pathway directions add important information when calculating production and degradation pathways, as they lend context to the interpretation of a gene expression change as it relates to the potential for production and degradation of different metabolites in the network.

### Calculation of production and degradation pathways for each metabolite

Then using this estimated flux state, we calculate weighted production and degradation pathways for each metabolite as follows. First, the reactions that carry flux in the estimated flux state are identified. Then, a desired pathway length D is defined. For each metabolite, reactions that are within the distance D by a forward traversal (in the case of degradation) or reverse traversal (in the case of production) of the flux carrying network are identified. For non-cofactor metabolites, cofactors were first removed from the network before traversal. The production or degradation pathway subnetworks are then extracted and mass balanced by adding compensating input and output reactions for unbalanced metabolites. These subnetworks are then broken down into elementary modes using a published algorithm(Chan and Ji 2011). Elementary modes are mass balanced pathways with weightings that when summed recapitulate the full flux distribution. Elementary mode pathways that contain the current metabolite are then extracted and summed to create a single weighted production or degradation pathway for the metabolite representing the contribution of reactions within a distance D to the production or degradation of the metabolite at the estimated flux state.

### Construction of aggregate pathway perturbation scores

To construct perturbed production and degradation scores for each metabolite, we first define reaction fold change scores by averaging the fold change for all genes that are involved in the catalysis of each reaction. We then define production and degradation pathway perturbation scores for each metabolite by calculating a weighted average of the pathways with their corresponding reaction expression fold changes. The weightings are assigned according to the reaction weightings within each pathway. These final production and degradation scores for each metabolite represent the expression change in reactions involved in the production and degradation of the metabolite, respectively, weighted by the degree of contribution of each reaction to the metabolite production/degradation at the estimated flux state.

## References

Bennett, Bryson D., Elizabeth H. Kimball, Melissa Gao, Robin Osterhout, Stephen J. Van Dien, and Joshua D. Rabinowitz. 2009. “Absolute Metabolite Concentrations and Implied Enzyme Active Site Occupancy in Escherichia Coli.” Nature Chemical Biology 5 (8): 593–99.

Bordbar, Aarash, Harish Nagarajan, Nathan E. Lewis, Haythem Latif, Ali Ebrahim, Stephen Federowicz, Jan Schellenberger, and Bernhard O. Palsson. 2014. “Minimal Metabolic Pathway Structure Is Consistent with Associated Biomolecular Interactions.” Molecular Systems Biology 10 (7). doi:10.15252/msb.20145243.

Chan, Siu Hung Joshua, and Ping Ji. 2011. “Decomposing Flux Distributions into Elementary Flux Modes in Genome-Scale Metabolic Networks.” Bioinformatics 27 (16): 2256–62.

Chubukov, Victor, Markus Uhr, Ludovic Le Chat, Roelco J. Kleijn, Matthieu Jules, Hannes Link, Stephane Aymerich, Jörg Stelling, and Uwe Sauer. 2013. “Transcriptional Regulation Is Insufficient to Explain Substrate-Induced Flux Changes in Bacillus Subtilis.” Molecular Systems Biology 9 (November): 709.

Duarte, Natalie C., Scott A. Becker, Neema Jamshidi, Ines Thiele, Monica L. Mo, Thuy D. Vo, Rohith Srivas, and Bernhard Ø. Palsson. 2007. “Global Reconstruction of the Human Metabolic Network Based on Genomic and Bibliomic Data.” Proceedings of the National Academy of Sciences 104 (6): 1777–82.

Fendt, Sarah-Maria, Joerg Martin Buescher, Florian Rudroff, Paola Picotti, Nicola Zamboni, and Uwe Sauer. 2010. “Tradeoff between Enzyme and Metabolite Efficiency Maintains Metabolic Homeostasis upon Perturbations in Enzyme Capacity.” Molecular Systems Biology 6 (April): 356.

Greenbaum, Dov, Christopher Colangelo, Kenneth Williams, and Mark Gerstein. 2003. “Comparing Protein Abundance and mRNA Expression Levels on a Genomic Scale.” Genome Biology 4 (9): 117.

Kanehisa, Minoru, Miho Furumichi, Mao Tanabe, Yoko Sato, and Kanae Morishima. 2017. “KEGG: New Perspectives on Genomes, Pathways, Diseases and Drugs.” Nucleic Acids Research 45 (D1): D353–61.

Khatri, Purvesh, Marina Sirota, and Atul J. Butte. 2012. “Ten Years of Pathway Analysis: Current Approaches and Outstanding Challenges.” PLoS Computational Biology 8 (2). Public Library of Science: e1002375.

Lake, Blue B., Rizi Ai, Gwendolyn E. Kaeser, Neeraj S. Salathia, Yun C. Yung, Rui Liu, Andre Wildberg, et al. 2016. “Neuronal Subtypes and Diversity Revealed by Single-Nucleus RNA Sequencing of the Human Brain.” Science 352 (6293). American Association for the Advancement of Science: 1586–90.

Orth, Jeffrey D., Tom M. Conrad, Jessica Na, Joshua A. Lerman, Hojung Nam, Adam M. Feist, and Bernhard Ø. Palsson. 2011. “A Comprehensive Genome-Scale Reconstruction of Escherichia Coli Metabolism--2011.” Molecular Systems Biology 7 (October): 535.

Orth, Jeffrey D., Ines Thiele, and Bernhard Ø. Palsson. 2010. “What Is Flux Balance Analysis?” Nature Biotechnology 28 (3): 245–48.

Subramanian, Aravind, Pablo Tamayo, Vamsi K. Mootha, Sayan Mukherjee, Benjamin L. Ebert, Michael A. Gillette, Amanda Paulovich, et al. 2005. “Gene Set Enrichment Analysis: A Knowledge-Based Approach for Interpreting Genome-Wide Expression Profiles.” Proceedings of the National Academy of Sciences 102 (43): 15545–50.

Wiback, Sharon J., Radhakrishnan Mahadevan, and Bernhard Ø. Palsson. 2003. “Reconstructing Metabolic Flux Vectors from Extreme Pathways: Defining the Alpha-Spectrum.” Journal of Theoretical Biology 224 (3): 313–24.

